# Unravelling vulture avoidance tactic of wind turbines combining empirical and simulation data

**DOI:** 10.1101/2023.07.26.550651

**Authors:** Sassi Yohan, Ziletti Noémie, Duriez Olivier, Robira Benjamin

**Author notes:** Corresponding author, CEFE, 1919 Routes de Mende, 34293 Montpellier, cedex 5, France. These authors contributed equally (Duriez Olivier, Robira Benjamin).

## Abstract

The increase of wind turbine installations to limit climate change may affect bird populations because of collisions with rotor blades. Birds may respond to wind turbine presence along a gradient of behavioural changes: avoiding the wind farm (macro-scale) or only the wind turbines either by anticipating wind turbine locations (meso-scale) or engaging into last-minute flee attempts after late perception (micro-scale). We investigated the flight response at these three spatial scales of 25 adult griffon vultures (*Gyps fulvus*) equipped with GPS tags over three years when flying in an area including ten wind farms in the Causses, France. At macro-scale, the population foraging range and habitat use revealed that vultures did not avoid wind farms. To investigate avoidance at meso- and micro-scales we focused on the four mostly visited wind farms. We compared vulture flights to null movement models, based on a method allowing us to keep the correlation between flights and topography while creating movement independent of wind turbine locations. At most sites, vultures did not show avoidance behaviour. Yet, simulations from our agent-based model highlighted that the avoidance pattern detected at one wind farm matched with an anticipated avoidance of turbines, probably linked to the presence of a ridge nearby. Overall, our results suggest wind farm-specific responses by soaring birds as a function of the landscape topography. Thus, stakeholders should carefully consider the wind farm location for siting and designing preventive measures (e.g. improve detection of species not able to avoid turbines in switching off on-demand technologies) to reduce collision risk of soaring birds.

## 1. Introduction

Wind turbines are a solution to produce electricity with limited CO_2_ emissions, although their impact on wildlife raise some concerns about the large-scale deployment of this technology. Meanwhile wind-power generation worldwide has grown dramatically during the last two decades (e.g. by 70% from 2015 to 2019; IPCC, 2022), mortality due to collisions with the rotor blades have been frequently reported in bats and birds (Schuster et al., 2015; Thaxter et al., 2017). Among birds, diurnal raptors are considered as one of the most vulnerable taxa (Thaxter et al., 2017) because of their slow pace of life which makes population viabilities particularly sensitive to additional adult mortality (Bellebaum et al., 2013; Carrete et al., 2009; Dahl et al., 2012; Duriez et al., 2022).

In response to wind turbine occurrence, birds can develop avoidance mechanisms at three spatial scales: macro-scale, meso-scale and micro-scale (May, 2015). Macro-scale avoidance refers to an avoidance of the wind farm as a whole (e.g. in Cabrera-Cruz & Villegas-Patraca, 2016; Plonczkier & Simms, 2012). Meso-scale avoidance describes an avoidance of the wind turbines several hundred to thousands of metres ahead (e.g. in Garvin et al., 2011; Santos et al., 2022; Schaub et al., 2020). Micro-scale avoidance stands for a last-second flee attempt of the rotor blades (typically < 200 m ahead) (May, 2015).

The avoidance tactic employed may be influenced by birds’ perception abilities, but also and largely by their morphology and flight capacities (Bevanger, 1998; Marques et al., 2014; Pennycuick, 2008). Several morphological parameters such as weight and wing area, which define wing loading, have been identified as determinants for collision risks (Janss, 2000). Birds with high wing loading, such as vultures and large eagles, have been shown to be more collision-prone than other raptors with lower wing loading such as common buzzard (*Buteo buteo*) or short-toed eagles (*Circaetus gallicus*) (Barrios & Rodríguez, 2004; de Lucas et al., 2008). The most likely reason for this pattern is that high wing-loading influences flight type (Shepard, 2022) and is associated with lower flight manoeuvrability (de Lucas et al., 2008). Unlike birds using flapping flight, large raptors use a soaring-gliding technique based on thermal and orographic updrafts to gain altitude effortlessly (Duriez et al., 2014; Shepard, 2022). Thermal and orographic updrafts, which are respectively masses of hot rising air emanating from heated surfaces and deviated wind onto topographical obstacles, constrain soaring birds in their displacement (Pennycuick, 1998). Thus, landscape features can also play an important role in the susceptibility of birds to collisions (de Lucas et al., 2012). While species with low wing loading could easily avoid wind turbines a few metres ahead (Schaub et al., 2020), those with high wing loading such as vultures will face much more difficulties. Despite being possible, a last minute flee attempt for large soaring birds requires them to switch to flapping flight, a flight mode they can not hold for long because of increased energetic costs (Duriez et al., 2014). Hence, if avoidance behaviour exists in these birds, we could expect an anticipated avoidance (meso-scale) allowing them to glide to their next updraft. This should particularly be true if the landscape favours thermal updraft or orographic uplift due to the surrounding topography.

Up to now, studies on wind turbines avoidance behaviours focused mainly on medium-sized birds with low wing loading such as black kites (*Milvus migrans;* Marques et al., 2020; Santos et al., 2022) or Montagu’s harrier (*Circus pygargus;* Schaub et al., 2020) which flight is relatively independent of landscape features. In this study, we adapted new methods to study avoidance behaviour in griffon vultures (*Gyps fulvus*), large soaring birds which depend largely on topography for their movements (Scacco et al., 2023). We investigated whether vultures actively avoided wind farms (macro-scale avoidance) and/or wind turbines (meso-/micro-scale avoidance). In the latter case, we aimed to characterise what was the flight response to wind turbines (i.e. progressive long-distance avoidance or last-minute flee attempt). Because of the dependence of their flight on the landscape, as well as their low flight manoeuvrability, we expected vultures to prioritise long-distance anticipated avoidance of wind turbines. Such in-depth investigations could particularly support stakeholder decisions by providing applied knowledge on where to site wind farms and how far to detect birds to shut down wind turbines in time to prevent collisions (McClure et al., 2021).

We used high-resolution GPS tracking of 25 adult individuals that ranged over 10 wind farms of the Causses region, France, over three years. To investigate macro-scale avoidance, we estimated vulture space utilisation distribution to determine whether vultures excluded wind farms from their ranging area. We coupled this with a habitat selection analysis to estimate in-flight selection of wind farms. To investigate meso- and micro-scale avoidance, we studied vulture movements within the four most intensively used wind farms and compared them to a null model of expected movements if independent of wind turbines location, obtained by rotating wind turbine locations. Furthermore, we compared true flights within wind farms to those simulated with an agent-based model to have a mechanistic understanding of the wind turbine avoidance manoeuvre (see Fig. 1 for framework).

**Fig. 1.**
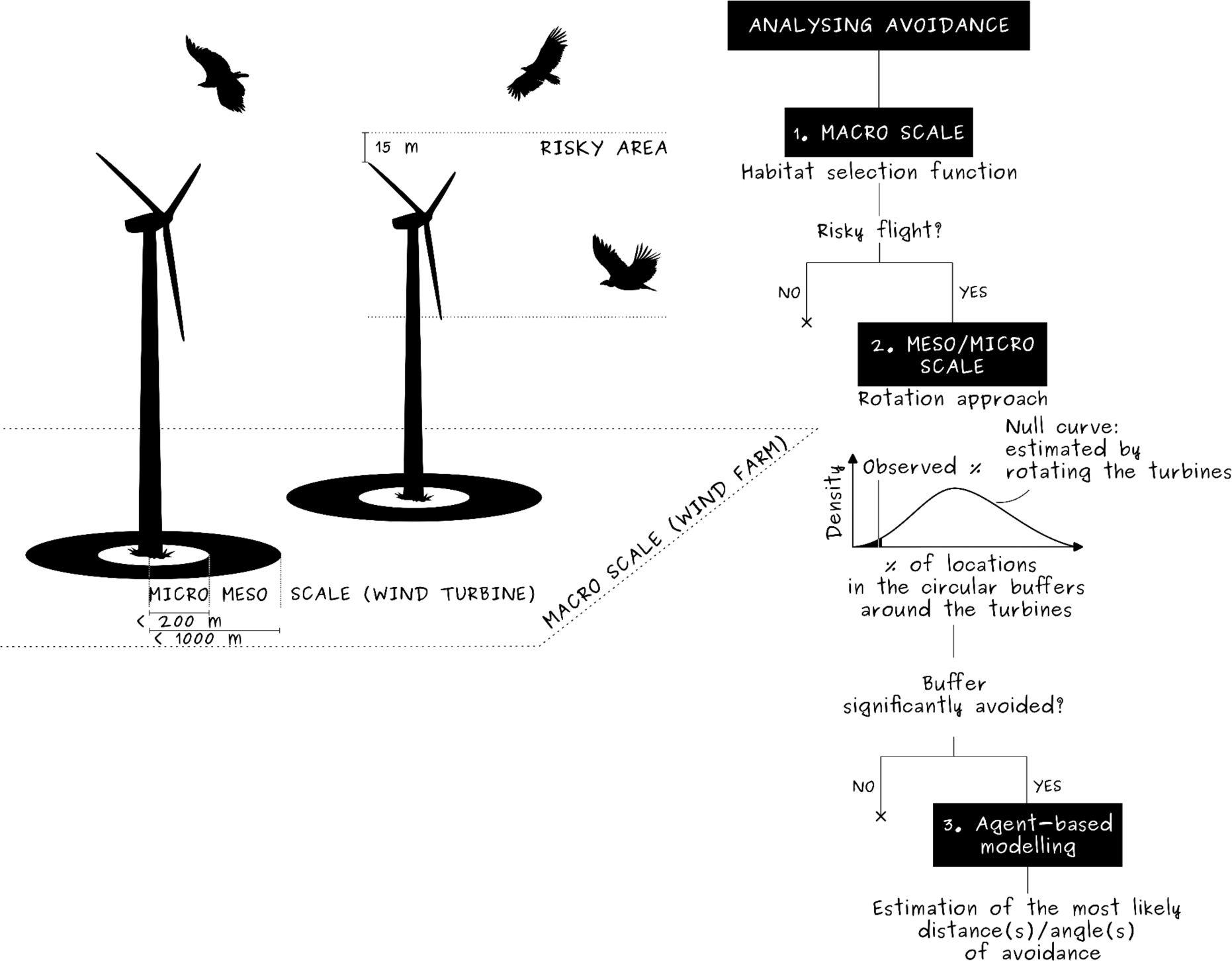
Illustration of the methodological framework used to investigate wind turbine avoidance in vultures. Avoidance tactics were studied at the macro scale (avoidance of the entire wind farm) or at the micro/meso scale (avoidance of wind turbines, from a hundred to a thousand metres) (left). We used a top-down approach from the largest to the smallest scale (right) combining empirical and simulation data.

## 2. Materials and methods

### 2.1. Study system

This study took place in the Causses region, France (Fig. 2), where a population of ca. 820 breeding pairs of griffon vultures live (census 2021, LPO). This region is characterised by limestone plateaux interspersed by valleys. Valleys offer conditions for orographic updrafts that vultures can use to soar efficiently. Away from the valleys, vultures patrol the open landscapes, relying on thermal updrafts to gain height, looking for mortality in herds of grazing livestocks. In recent years, both the number of vultures and the number of wind turbines have increased. There are nowadays 10 operating wind farms (totalling 130 turbines) and nine additional are planned (projects totalling 91 turbines, Fig. 2), in a region where at least 30 vultures have been found dead due to collisions between 2012 and 2022 (including 10 casualties at the four focal wind farms cited below) (LPO/DREAL Occitanie, unpublished). These wind farms are located between 18 km and 52 km from Cassagne, the geographical centre of the breeding colony where a collective natural recycling station with vultures stands (44°12’N, 3°15’E, Fig. 2, Duriez et al., 2021).

**Fig. 2.**
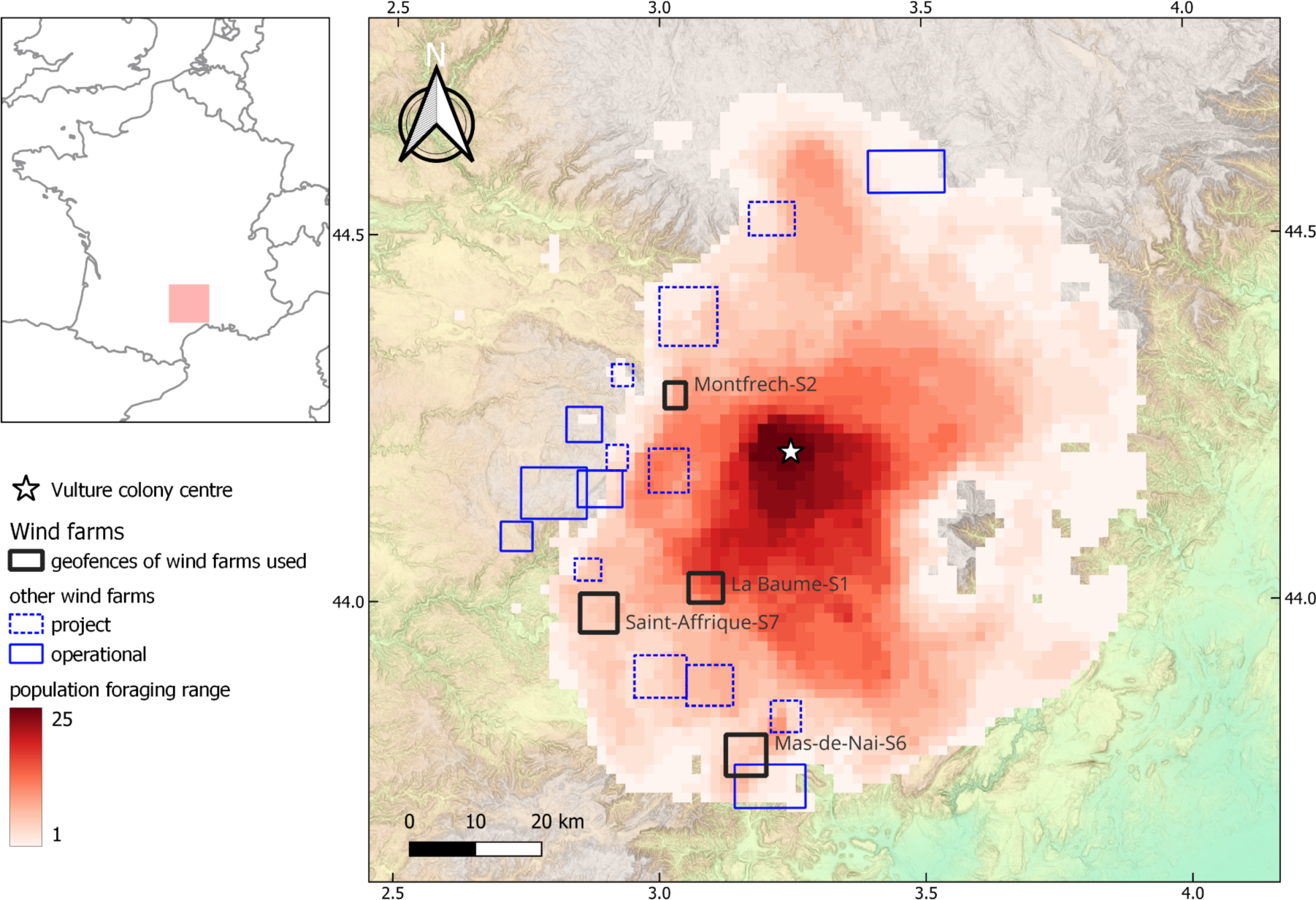
Global vulture population foraging range in the Causses region, France. The darker the colour of the red raster, the higher the number of individual in-flight 95% utilisation distributions that overlap the given cell. Additional rectangles represent geofences around wind farms (black: study wind farms, plain blue: operational wind farms; dotted blue: project of construction). The star is the geographical centre of the nesting colony, Cassagne, where vultures have been reintroduced, are tagged and where a carcasse recycling station is located.

We used tracking data spanning 3 years (from 1^st^ January 2019 to 31^st^ December 2021) from 25 vultures (Table S1) that had been captured in 2018 at Cassagne carcasses recycling station, and equipped with 50 g solar-powered GPS-GSM tags (Ornitrack-50, Ornitela), in a leg-loop harness configuration (Anderson et al., 2020). GPS tags were set to record location, speed and altitude at intervals of 2-15 min depending on battery levels and season (generally lower battery levels in winter). To study avoidance behaviour of operating wind turbines by vultures, we defined rectangular geofences (virtual barriers) placed at 2 km from the most outlying turbines in each wind farm. Within these geofences, GPS tags automatically shifted to high resolution recording (1 Hz) of individuals’ location, speed and altitude. This 2 km threshold was defined based on a compromise between the need of time for the tags to switch to high resolution before entering the 1 km meso-scale buffer, and the need to prevent battery discharge by recording at high resolution in areas that we were not interested in. To retain only accurate in-flight GPS locations, we filtered the GPS locations of each individual by their groundspeed (> 4 m/s) and their horizontal dilution of precision index (HDOP < 4) (Martin-Díaz et al., 2020; Nathan et al., 2012). Data cleaning, processing and analysis were performed with *R* (version 4.2.2, R Core Team, 2022).

### 2.2 Data analysis

#### 2.2.1. Macro-scale avoidance

To find out whether vultures expressed a macro-scale avoidance of wind farms, we computed an in-flight utilisation distribution (UD) and an habitat selection function for each individual. First, we resampled flights every 10 min to homogenise the sampling frequencies (“track_resample” function, *amt* R package, Signer et al., 2019). Then, we focused on movements that were at a distance < 55 km of the colony centre. This distance enabled the inclusion of all wind farms of the region while focusing on vultures’ daily flights (mean daily displacement from Cassagne by local birds equals 26 km (SD ± 10 km), Fig. S1, Fluhr et al., 2021). Individuals’ UDs were estimated on these flights using brownian random bridge-based kernels (Benhamou, 2011, *adehabitatHR* R package (Calenge, 2006), see supplementary materials ESM01 for details). We then estimated a “population foraging range” as the layering of the 95% isopleth of individual UDs where each cell value corresponded to the number of individual UDs overlapping that cell (Duriez et al., 2019).

To estimate if vultures tended to fly further from wind turbines than expected by chance we computed an habitat selection function (HSF; Fieberg et al., 2021). To do so, for each individual we subsampled its daily datasets at three locations per day, evenly spaced during the main activity period of vultures and not temporally autocorrelated (at 10:00, 12:00, 14:00; Fluhr et al., 2021). This allowed us to categorise the locations “used” by individuals. In parallel, as the tracked vultures are central place foragers (Monsarrat et al., 2013), we sampled 10-fold more locations following a bivariate exponential distribution (“available locations”, Benhamou & Courbin, 2023). We restricted these locations within a distance of 55 km from the colony centre. We fitted an HSF for each individual, using the distance to the closest operational wind turbine as the only predictor. Each HSF corresponded to a generalised linear mixed model with a binomial error structure (available: 0, used: 1) and a weighted logit link function considering a weight of 5000 for available locations, and 1 for used locations. The exponential of the unique slope estimate indicates whether vultures show no preference (≈ 1), favour wind farms (> 1) or avoid wind farms (< 1) (Fieberg et al., 2021).

#### 2.2.2 Meso- and micro-scale avoidance

To investigate meso- and micro-scale avoidance behaviour we focused on four wind farms: La Baume, Montfrech, Mas de Naï and Saint Affrique. These wind farms were among the closest to the centre of the vultures’ colony and were the most visited ones by vultures (Fig. 2, Table S2). Among the 25 vultures, 92% of them crossed at least once one of these four operating wind farms within the rotor swept zone during the three years considered (Table S1).

##### 2.2.2.1 Use of topography within wind farm geofences

In the geofenced areas of these winds farms, orographic updrafts are generated by steep slopes associated to valleys, which are easily identifiable by a human eye in the coloured topography rasters presented in Fig. 3 (Digital Elevation Model, IGN BDTOPO, 25 m resolution). Hence, we estimated the central value of elevation among all pixels of the raster (i.e. 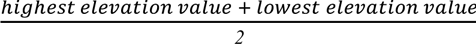 and we created an isoline of elevation at this value. Then, because orographic updrafts are generally drifted towards the upper part of the ridge we empirically used a 300 m buffer to geographically define the area (hereafter called “slopes”) most likely to generate orographic updrafts. To estimate how topography constrained vulture flight we computed another HSF. Here we empirically found that subsampling 30% of the GPS locations composing each vulture track in the considered geofenced area gave robust results while reducing autocorrelation between locations. The locations “available” to vultures were randomly sampled within the geofenced area. The HSF used to estimate the preference for slopes over other areas followed the same structure as mentioned above with a dummy variable indicating whether the location was within a slope (1) or not (0) as a unique predictor.

**Fig. 3.**
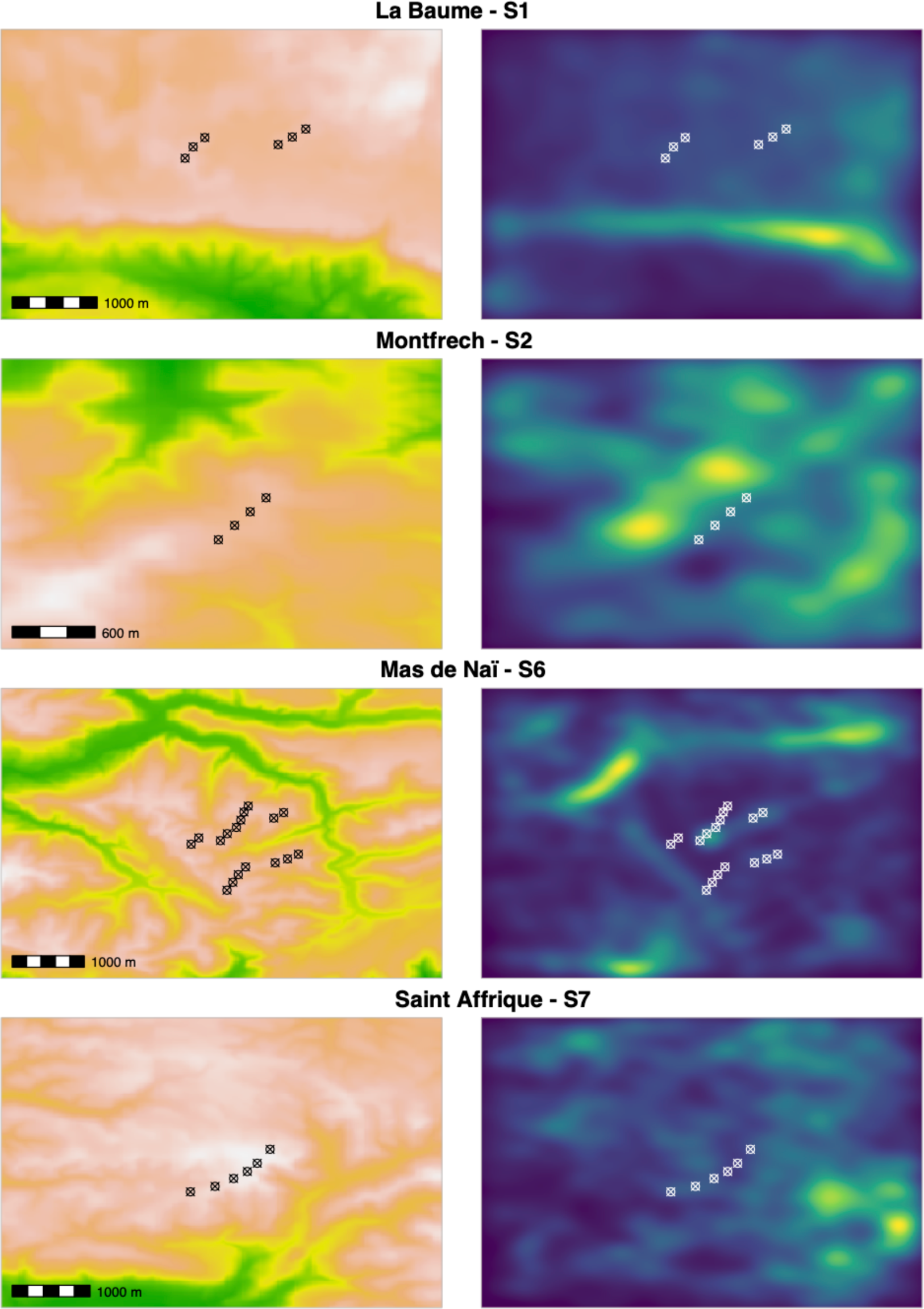
Topography and vulture space utilisation distribution in the studied wind farms. Topography distribution (left column; green to white gradient indicating increasing altitude) and the weighted mean utilisation distribution (right column, blue to green gradient indicating increasing use intensity). See supplementary materials ESM01 for details on the estimation of the mean utilisation distribution.

##### 2.2.2.2 Wind farm rotation to create null model

To investigate whether vultures anticipated wind turbine locations to start manoeuvring at long distance and/or whether they performed short-distance reflex manoeuvre to avoid them, we separated flights for which vultures flew within the rotor-swept zone of wind turbines (i.e. between the minimum rotor tip height and 15 m above the maximum rotor tip height, see Table S2 for wind farms’ specific values) and those for which vultures flew above the rotor-swept zone. These flights were rediscretised at constant step length (50 m), to remove bias due to speed differences within and between the tracks but also to reduce location aggregation due to circular soaring phases compared to rectilinear gliding phases. Vultures can also fly below the rotor swept zone, yet, these events are rare (3.5 % of the locations are below the rotor swept zone in our study), thus we did not consider that case.

We defined avoidance behaviour as a use of an area containing wind turbines lower than expected if vulture flew independently of the wind turbine positions. To do so, we first estimated the percentage of locations occuring within a given range of turbines (buffer zone) at their original positions (e.g. 8.92% of the observed locations are within a 300 m buffer around wind turbines in the example shown in Fig. 4A). Then we compared this observed percentage to a null distribution expected if turbines were not avoided. We created this null distribution by recalculating the percentage of locations included into the same buffer zone when the geofenced area containing the wind farm was rotated around its barycenter from 10° to 350° with a 10° step (e.g. 12.33% of locations were included into the 300 m buffer with a 10° rotation in the example shown in Fig. 4B). Rotating the wind farm, instead of the flight tracks, allowed us to preserve correlations between flights and topography, a necessary condition for the null model to be biologically meaningful (Martin et al., 2008). This process, repeated over buffers ranging from 50 to 1000 m from each turbine (with 50 m steps), provided a null distribution associated with each buffer size.

**Fig. 4.**
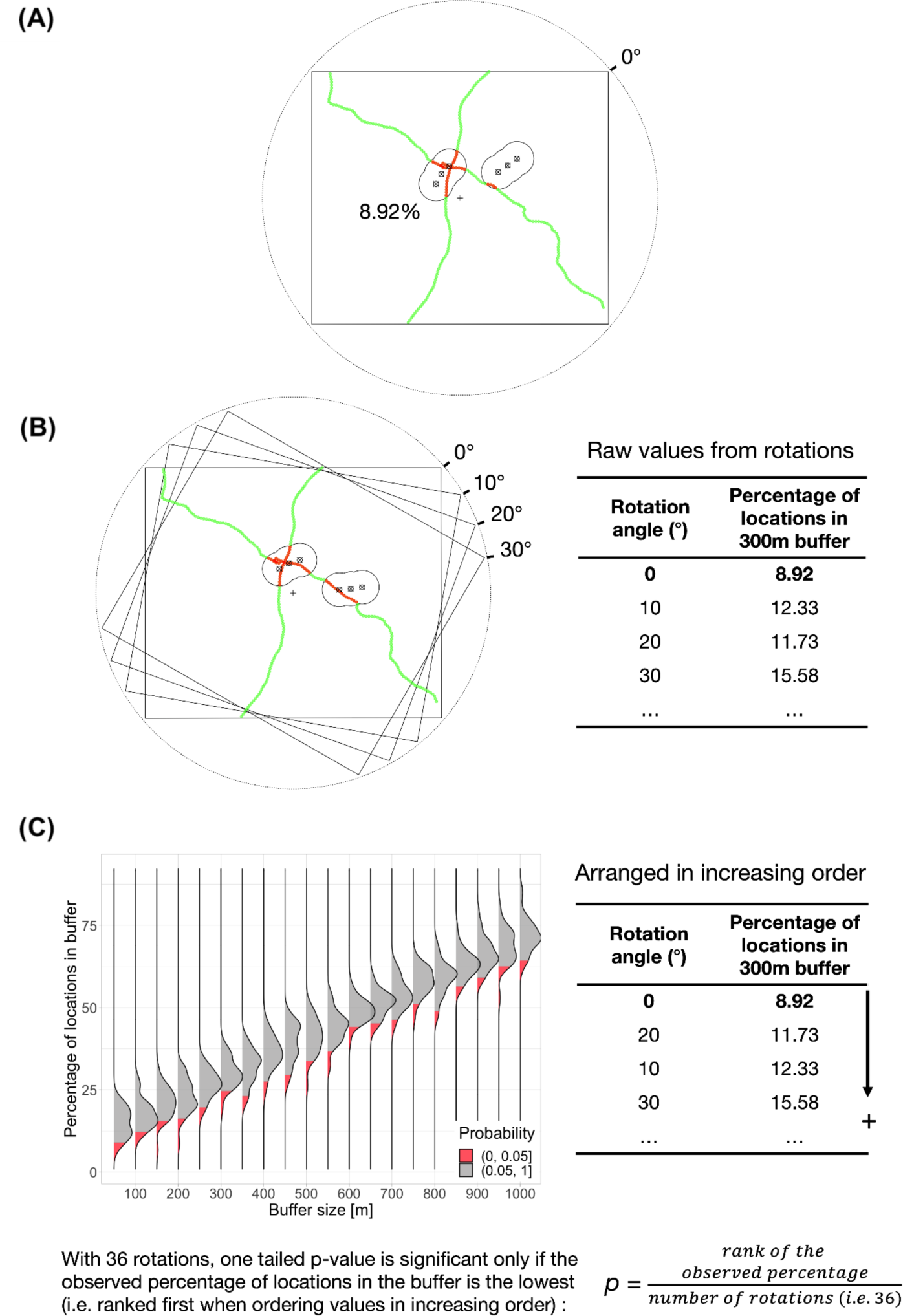
Visual guide of the rotational approach used to estimate vulture turbine avoidance. (A) The geofence is represented by the rectangle around the wind farm (e.g. La Baume here). Wind turbines are indicated by the crossed circles and are surrounded by a specific buffer zone (e.g. 300 m here), the merged limits of which are shown by the solid black line. Two vulture flights are represented by the green and red lines (outside or inside buffers, respectively). For the original (true) wind turbine positions (0° of rotation), the percentage of vulture locations observed in the buffer was estimated. (B) A null model was constructed by rotating the wind turbines around the wind farm barycentre from 10° to 350° by steps of 10°. For each rotation the percentage of vulture locations in the considered buffer was calculated, giving 36 values of percentage per buffer size (1 observed and 35 theoretical values). (C) This method was applied to buffers from 50 m to 1000 m by steps of 50 m around wind turbines, providing a null distribution of expected percentage for each buffer size. An avoidance should be detected if the truly observed percentage of locations in the buffer (i.e. for rotation angle = 0°) fell into the red part of the distribution. Hence, for each buffer size the percentages of locations were ranked by increasing order and we estimated the one-tailed p-value by dividing the rank of the truly observed percentage in the arranged distribution by the total number of rotations. Thus, the observed percentage of locations in the buffer was significantly lower than expected if it was ranked first in the distribution: *p* = 1/36 = 0.028. (These are simulated data for illustration)

For each buffer, we defined it as avoided when the observed proportion of locations was significantly lower than expected through the null distribution. For this purpose, we ranked, by increasing order, the 36 percentage values obtained for the considered buffer (the observed value, at rotation = 0°, plus the 35 values from the rotations). We estimated the one-tailed probability (hereafter *p*) by dividing the observed proportion rank by the total number of values (i.e. 36; Fig. 4C). A significant avoidance of the buffer was detected (*p* < 0.05) when the observed percentage was ranked as the lowest (i.e. *p* = 1/36 = 0.028). We applied this procedure for both flights within and above the rotor swept zone separately. We checked that this rotation approach could adequately identify avoided buffers in each wind farm when simulating different avoidance scenarii with our agent-based model described in the following section (supplementary materials ESM02 and Fig. S2).

##### 2.2.2.3 Agent-based model simulations

When a significant avoidance pattern was identified with the above-mentioned procedure, we aimed at determining whether this pattern fitted with a long-distance anticipated avoidance or a last-minute flee attempt. We built an agent-based model (DeAngelis & Mooij, 2005; Grimm & Railsback, 2005) simulating the behaviour of a virtual vulture able to perceive a turbine (and start manoeuvring) at a distance *d*, and able to adjust its heading (known as turning angle) of *α*° every 5 m. This 5 m step was meant to be as small as possible to mimic continuous movement, but sufficiently reasonable due to computational limits. The environment, in which a virtual vulture flew, contained wind turbines whose positions matched the true configuration of the studied wind farms.

Each simulation followed the subsequent flow:

1. A starting location was randomly selected on a side of the considered geofenced area and the target location (reached only through strict ballistic movement) was defined on the mirroring side, such that the target direction was *θ*.
2. The virtual vulture started moving in the direction of the target location, following a biased random walk (Codling et al., 2008). This biased random walk consisted of movement steps of 5 m in the direction *θ’* sampled in a Von Mises distribution of mean *θ* (“rvm” function of the *CircularDDM* package; Lin et al., 2018), and of persistence value *κ* (estimated based on the true vulture tracks occurring at wind farms of comparison, rediscretised at a 5 m interval, using the “est.kappa” function of the *CircStats* package, Lund & Agostinelli, 2018).
3. If the virtual vulture arrived at a defined distance of *d* metres from a wind turbine, it engaged in an avoidance behaviour which consisted in maintaining a turning angle of *α*° opposite to the turbine location (e. g. left if the turbine was initially located right with respect to the heading at start of avoidance, and *vice versa*) at each step.
4. If the virtual vulture was avoiding a turbine, avoidance behaviour stopped as soon as the distance to the turbine started to increase. It then resumed its biased random walk (heading to the last direction after avoidance), and would return into avoidance behaviour whenever a new turbine was perceived.
5. The simulation stopped when the agent flew out of the geofenced area.

We tested jointly for several values of *α* (0° to 14° by steps of 1°; 14° representing the maximum angle a griffon vulture can turn within a thermal; Williams et al., 2018) and *d* (50 m to 1000 m by a step of 50 m, similar to the rotation procedure), repeating the simulations 10 000 for each set of parameters. A small *α* and a large *d* would mimic long-distance anticipated avoidance, while a large *α* and a small *d* would mimic last-minute flee attempt.

To understand the movements rules underpinning vultures’ avoidance behaviour, we focused on buffer sizes detected as avoided, and compared the absolute fit of the simulations with the empirical data. The absolute fit corresponded to the square of the difference between the percentage of locations obtained by simulations and the one observed on empirical data (ϑ). In simulations mimicking avoidance (*d* and *α* > 0), we removed the ones for which the fit with empirical data was worse than for cases with no avoidance (ϑ/ϑ_null_ < 1 where ϑ_null_ is the percent of location obtained in the buffer when simulating no avoidance). For the remaining cases, we defined the fit quality between empirical and simulated percentage of location with a buffer as *f* = 1 - ϑ/ϑ_null_. A perfect fit (i.e. the combination of *α* and *d* that correctly mimicked the observed vulture avoidance behaviour in the considered buffer) would give *f* = 1.

## 3. Results

### 3.1 Avoidance behaviour of wind farms at macro-scale

The population foraging range overlapped with 100% of the wind farm projects and 60% of the operating wind farms (Fig. 2). The turbines of the furthest wind farm from the colony centre, Mas de Naï, was included into the in-flight utilisation range of five vultures while one of the closest wind farm, La Baume, cut off the airspace used by 18 individuals (Fig. 2). In addition, the mean exponential of the HSF estimate associated with the closest distance to operational wind turbines was extremely close to 1 (0.99 ± SD 4.82 × 10^-5^) suggesting no preferences toward large distances from wind turbines.

### 3.2 Avoidance behaviour of wind turbines at meso-/micro-scale

#### 3.2.1 Importance of the topography when flying in the wind farms

Vultures crossed the wind farms several times during these three years (all individuals pooled, [min,max] = [207, 1793] in Montfrech and La Baume, respectively - Table S1 and S2). The proportions of tracks that entered the rotor swept zone of these wind farms were not negligible ([min,max] = [32.84%, 50.28%] in Montfrech and Mas de Naï, respectively).

In the area defined by the geofences, 15.3% of La Baume, 51.5% of Montfrech, 80.5% of Mas de Naï and 19.7% of Saint Affrique, were represented by slopes (Fig. 3). While flying in these areas, vultures significantly favoured these slopes (exponential of HSF estimate associated to slope use [95% confidence interval]; La Baume: 1.807 [1.767,1.848], p < 0.001; Montfrech: 1.193 [1.106,1.287], p < 0.001; Mas de Naï: 1.306 [1.246,1.370], p < 0.001; Saint Affrique: 1.143 [1.088,1.200], p < 0.001).

#### 3.2.2 Detection of active avoidance of wind turbines

In three wind farms (La Baume, Mas de Naï, and Saint Affrique), no avoidance behaviour was detected either for flights above or within the rotor swept zone (Fig. S3). In Montfrech, we did not detect avoidance when focusing on flights above the rotor swept zone (Fig. 5A). However, for flights within this risky zone, we observed a significantly lower proportion of GPS locations than expected for buffers from 50 m to 450 m (Fig. 5B). This suggests a significant avoidance in this range of distances, matching with the availability of steep slopes nearby (Fig. 5C).

**Fig. 5.**
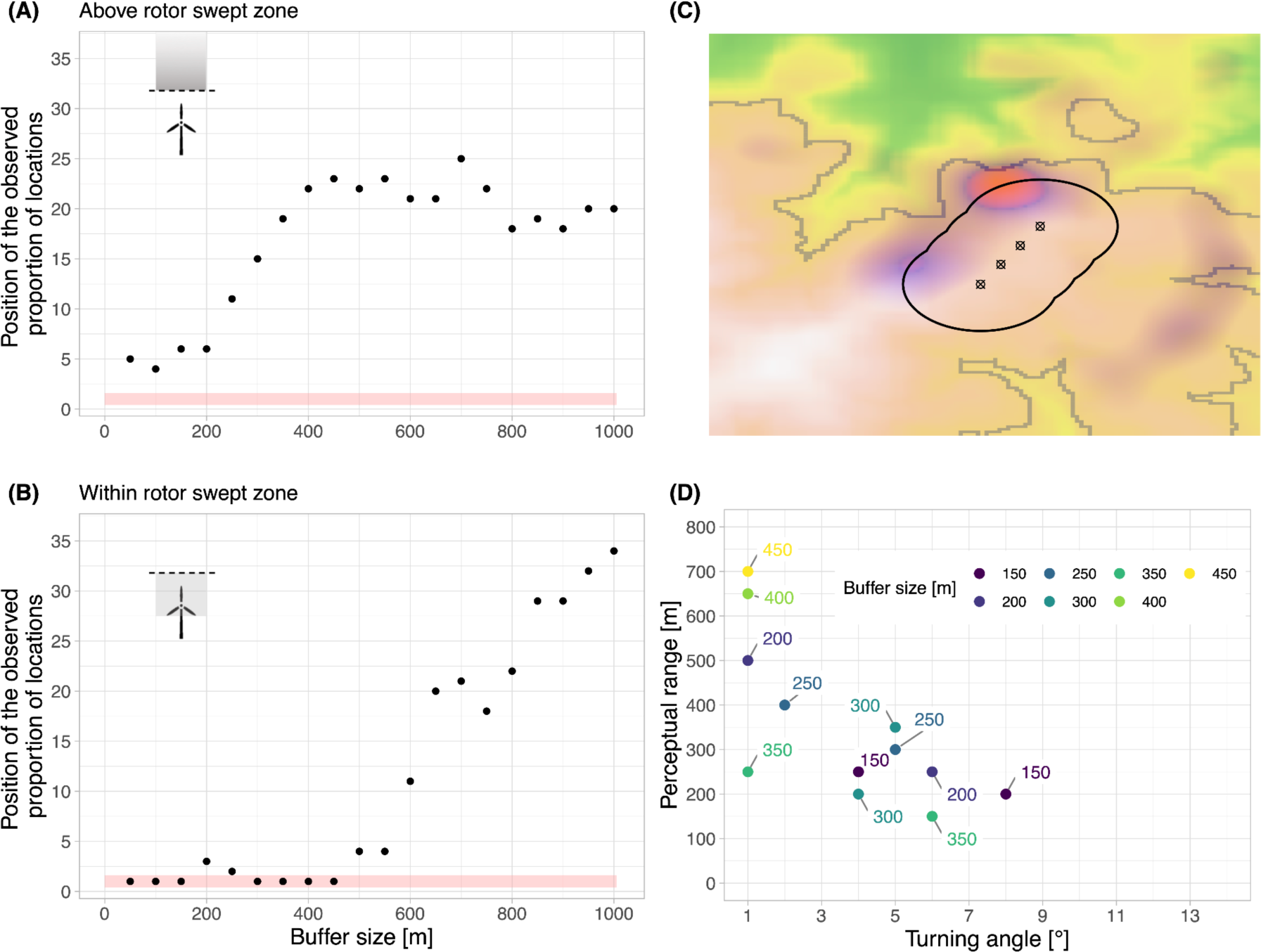
Turbines avoidance in Montfrech wind farm. Results of the rotation analysis conducted on vultures’ flights above (A) and within (B) the rotor swept zone: dots represent the position of the observed percentage of location, among the 36 estimated values (*y-*axis) during the rotational analysis, within a given circular buffer around wind turbines (*x*-axis). The red rectangle highlights buffers for which the amount of locations observed is significantly lower than randomly expected. (C) illustrates the overlap between topography (green to white gradient indicating increasing altitude) and utilisation distribution (purple to red gradient indicating increasing use intensity) of the Montfrech wind farm. The solid black line represents the limits of merged 450-m buffers around wind turbines. (D) shows the combination of parameters (perceptual range and turning angle) used in the agent-based model to simulate vulture flights yielding the best fits between simulations and observations (*f* > 0.95).

#### 3.2.3 Characterisation of meso- and micro-scale avoidance tactic

We further scrutinised flights in Montfrech, by comparing the amount of vulture locations observed within buffers around wind turbines, where avoidance was detected, to the amount obtained with agent-based simulations. Simulations with large distances of detection and low turning angles yielded best matches with observations for most of the buffers highlighting a predominant long-distance avoidance tactic (Fig. 5D). However, for the smallest buffers the best fit (*f* > 0.95) between simulated and observed data could be obtained for both low and large turning angle values, suggesting potential last-minute flee attempts (e.g. the observed avoidance of the 200 m buffer is well simulated with both sets of parameters: *d* = 500 m with *α* = 1° and *d* = 250 m with *α* = 6°, Fig. 5D).

## 4. Discussion

Combining high-resolution GPS tracking data from 25 adult griffon vultures across four wind farms and simulations from an agent-based model revealed that wind turbine avoidance seems relatively limited in this species, possibly with specific responses in each wind farm strongly associated with landscape topography. The landscape surrounding wind farms thus appears of prime importance and should thus prevail for deciding wind farm sitting.

Our exploration of the population foraging range coupled with the habitat selection analysis highlighted that wind farm areas were still exploited by vultures. Precisely, we demonstrated that griffon vultures did not exhibit a macro-scale avoidance of the wind farms. This reminds of former results on other large raptors such as white-tailed eagles (*Haliaeetus albicilla*; Dahl et al., 2013) but contradicts results of macro-scale avoidance in medium-size birds such as migrating raptors at onshore wind farms (Cabrera-Cruz & Villegas-Patraca, 2016) and aquatic birds at offshore wind farms (e.g. in Plonczkier & Simms, 2012).

At meso-scale, we observed signs of avoidance behaviour, up to 450 m, only at one wind farm (Montfrech) among four commonly crossed by vultures. Unlike other wind farms, Montfrech turbines were located at about 500 m from steep slopes that were significantly selected by vultures for their foraging and commuting movements. Such a site-specific response could be explained by the topography of Montfrech, where vultures could reach slopes to take advantage of orographic uplift and fly parallel to the row of turbines (like black kites in Santos et al., 2022). Since it only requires positioning themselves over windward slopes to benefit from the deviated wind above canyons’ ridges and slopes, the predictability of orographic updrafts make them easier to exploit compared to thermals (Katzner et al., 2012; Shepard, 2022). As such, the avoidance of wind turbines identified by our analyses could be “passive”, as a by-product of topography, rather than an active avoidance due to a perceived threat. Indeed, as the strength of the slope uplift decreases with height above ground level (Shepard, 2022), vultures flying above 200 m over ground would no longer be able to rely on this source of uplift (Duerr et al., 2019). This framed consistently with the lack of avoidance when considering flights above the rotor swept zone at all wind farms, including Montfrech.

Far away from used slopes and ridges, as in the three remaining wind farms where we did not detect any avoidance, soaring birds may rely almost exclusively on thermals to gain altitude (Katzner et al., 2012). Péron et al. (2017) estimated that the probability for griffon vultures and other large raptors to fly above 200 m (i.e. above the rotor-swept zone) was significantly correlated to thermal uplift potential. Being constrained in their movements by such unpredictable resources may explain why soaring bird mortality by collision on wind turbines increases when thermals are less frequent or less powerful (e.g. during rainfall or during winter; Barrios & Rodríguez, 2004; Marques et al., 2014). The circling flight necessary to rise into thermal may be associated with a higher risk of collision than when using (linear) slope soaring due to repeated passages in the same area at increasing altitudes (Barrios & Rodríguez, 2004). This may also explain why many bird species that rely on the same flight tactics suffer heavy losses by collisions (Barrios & Rodríguez, 2004; de Lucas et al., 2008; Heuck et al., 2019; Katzner et al., 2012) and concur with a fairly low support for a last-minute flee attempt in our analyses.

Such last-minute avoidance should require sharp turns, which are better achieved by birds with low wing-loading and elongated tails, which is not the case of *Gyps* vultures that possess a rather short tail and high wing loading (Balmford, 1995; Gillies et al., 2011). Such manoeuvre could still occur in rare cases of emergency but would result in a very rapid loss of altitude for the vultures. This picture, however, contrasts partially with recent evidence on black kites and Montagu’s harriers showing that flight behaviour was modified at a close range of wind turbines suggesting an active avoidance in these species (Santos et al., 2022; Schaub et al., 2020). These species, having a lower wind loading, are probably much less constrained by sources of uplift in their movements, giving them more room to adapt their flights according to perceived risks, likely explaining the observed differences in avoidance behaviour.

In this study we adapted a method consisting of rotating the locations of infrastructure to be avoided (here wind turbines) to construct null models distributions of space use independent of infrastructure locations. This method is similar in principle to the usual method to rotate tracks to create a null model (e.g. in Schaub et al., 2020) but it has the double advantage of saving the correlation between topography and animal flight, and also reducing the computation power (and time) needed to perform rotations of large amounts of tracks. This is particularly valuable for species heavily relying on topography to move through landscapes, whose rotated flight would become biologically unrealistic. In addition, the use of an agent-based model allowed us to highlight the robustness of our method and be confident in the pattern detected on empirical data. It is also a practical approach for understanding the processes underlying animal movements (Tang & Bennett, 2010), but is often overlooked for analysing collision and turbine avoidance by flying animals (but see in birds: Eichhorn et al., 2012, in bats: Ferreira et al., 2015). We have provided here a model that can be used as a framework for further investigation of the risk of collision with wind turbines. All together, our results revealed a new level of complexity in wind turbine avoidance behaviours as even among restricted groups such as soaring raptors, answer to turbine presence seem to be species- and site-specific.

## 5. Conclusion and management implications

The tragic conflict that we currently face is that soaring birds and wind energy developers are targeting the same resource: wind. The development of wind farms pose a major conservation problem for most large flying animals (Thaxter et al., 2017), as they can induce disturbance of the environment, leading to a decrease in local biodiversity, and can also lead to disruptions of population dynamics and stability through collision fatalities (Perrow, 2017). Here, we provided further evidence that the flight capabilities of some species may make them particularly sensitive to wind turbine collisions, and do not allow them to avoid wind turbines effectively. Yet, we detected an anticipated avoidance at one wind farm matching with the presence of slopes. Slopes aggregate soaring birds and may allow them to stay away from turbines, provided they are neither too close (high risk of collision using the slope uplift) nor too far away (high risk of collision using only thermal uplift; Péron et al., 2017). Taking into account distance from turbines to favourable conditions when sitting projects could help to reduce collision risk. However this would require further research to first understand what makes some slopes more attractive to soaring birds, as all slopes are not necessarily used. Furthermore, it would imply a better understanding of the distance which would be optimal to reduce collision risk. At already operating sites it has become crucial to detect birds unable to avoid turbines well in advance to shutdown turbines in time to prevent collisions. Shutdown on-demand when animals at risk are detected is a potentially promising way to reduce collision mortality with a negligible reduction in energy production, yet automatic detection systems are costly and their efficiency is still debated (McClure et al., 2021; Tomé et al., 2017). Straightforwardly, to solve this green-green dilemma to reduce carbon emission and preserve biodiversity, it would be more efficient, and should be prioritised, to prevent siting the turbines at places where soaring birds are obliged to travel.

## CRediT authorship contribution statement

**Yohan Sassi**: Conceptualization, Methodology, Software, Formal analysis, Visualisation, Writing - Original Draft, Writing - Review & Editing. **Noémie Ziletti**: Conceptualization, Funding acquisition, Writing - Review & Editing. **Olivier Duriez**: Conceptualization, Data curation, Visualisation, Writing - Review & Editing, Supervision. **Benjamin Robira**: Conceptualization, Methodology, Software, Formal analysis, Visualisation, Writing - Original Draft, Writing - Review & Editing, Supervision.

## Data availability

GPS telemetry data are stored in the www.movebank.org database in the study “Eurasian Griffon vulture in France (Grands Causses 2018) ID_PROG 961”. Given the sensitive nature of tracking data for protected species, downloading permission should be asked through the platform to OD. Scripts for review are available here: https://github.com/YohanSassi/windTurbinesAvoidance. A perennial storage will be provided after revision (e.g. Zenodo).

## Acknowledgment

We thank the staff from the Grands Causses site of LPO France (T David, R Straughan, R Nadal L Giraud, P Lecuyer, B Descaves) for helping in capturing vultures for tagging. Telemetry study of vultures was authorised in the Programme Personnel 961, coordinated by O. Duriez, under the supervision of the French ringing centre, CRBPO, Paris. We also thank Patrick Boudarel (DREAL Occitanie, France) for fatalities data in the Occitanie region, France and Simon Benhamou for helpful discussions regarding data analyses.

