## Supplementary materials for "Unravelling vulture avoidance tactic of wind turbines combining empirical and simulation data"

1: CEFE, Univ Montpellier, CNRS, EPHE, IRD, Montpellier, France

2: LPO France site Grands Causses, Le Bourg, Peyreleau, France

3: Department of Biodiversity and Molecular Ecology, Research and Innovation Centre, Fondazione Edmund Mach, San Michele all'Adige, Italy

† These authors contributed equally.

### Supplementary materials

ESM01: Additional methods for utilisation distribution

To explore the space used by vultures at macro-scale we computed an in-flight utilisation distribution (UD). UDs were estimated using brownian random bridge-based kernels (Benhamou, 2011) using the ”BRB.D” function (to first estimate the coefficient of diffusion) and the “BRB" function (to estimate the UD) of the *adehabitatHR* R package (Calenge, 2006). Because of vultures’ mean flight speed of 50 km/h (Williams et al., 2018), we considered locations as independent (i.e. no inference of a bridge) when the time interval exceeded 1 h (parameter $T_{max}$). We considered static phases when vultures travelled a distance < 20 m during the 10 min interval (i.e. higher than GPS precision, parameter $L_{min}$) and used a smoothing parameter ($h_{min}$) of 500 m.

To explore the space used by vultures in wind farm geofences, we used the same method as described above but on rediscretised individual flights (with a constant step of 50 m). We changed the parameters as follows. We considered locations as independent (i.e. no inference of a bridge) when the time interval exceeded 30 s (parameter $T_{max}$) due to the high resolution tracking. We considered static phases when vultures travelled a distance < 20 m (i.e. higher than GPS precision, parameter $L_{min}$) and used a smoothing parameter ($h_{min}$) of 150 m. As the number of flights in the wind farms varied greatly between vultures (Table S1), the population UD was obtained here by previously weighting each individual 95% UD by the number of flights in the wind farm of interest.

ESM02: additional methods to assess validity of the rotational approach

The rotational procedure identified avoidance patterns accurately when we simulated 1000 tracks, each time with the following combination of parameters: a perceptual range ($d$) of 450 m and a turning angle ($\alpha$) of 0°, 1° or 5° for the three first scenarios, respectively and $d$ = 50 m with $\alpha$ = 5° for the last scenario. Specifically, for the two first scenarios in Montfrech, it did not detect any avoidance when $\alpha$ = 0°, but as soon as we simulated an avoidance with $\alpha$ = 1°, it accurately detected a significantly lower proportion of locations in buffers up to 450 m.

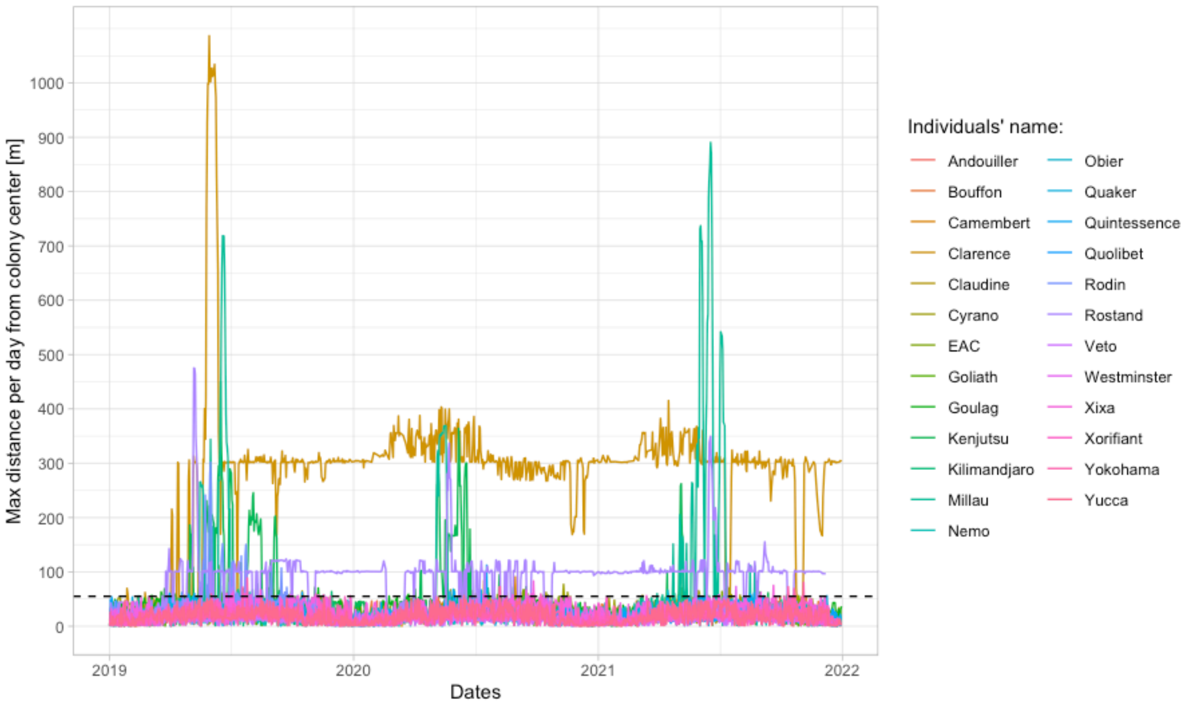


**Fig. S1. Maximum daily distances from the centre of the colony reached by vultures between 2019 and 2021**. A threshold of 55 km from the centre of the colony is highlighted by the horizontal black dotted line. Two individuals, Clarence and Rostand, mostly settled in Pyrenees and Ardeche respectively, > 100 km from the Causses, France and were discarded in the estimation of the mean maximum displacement from the centre colony reached daily by vultures.

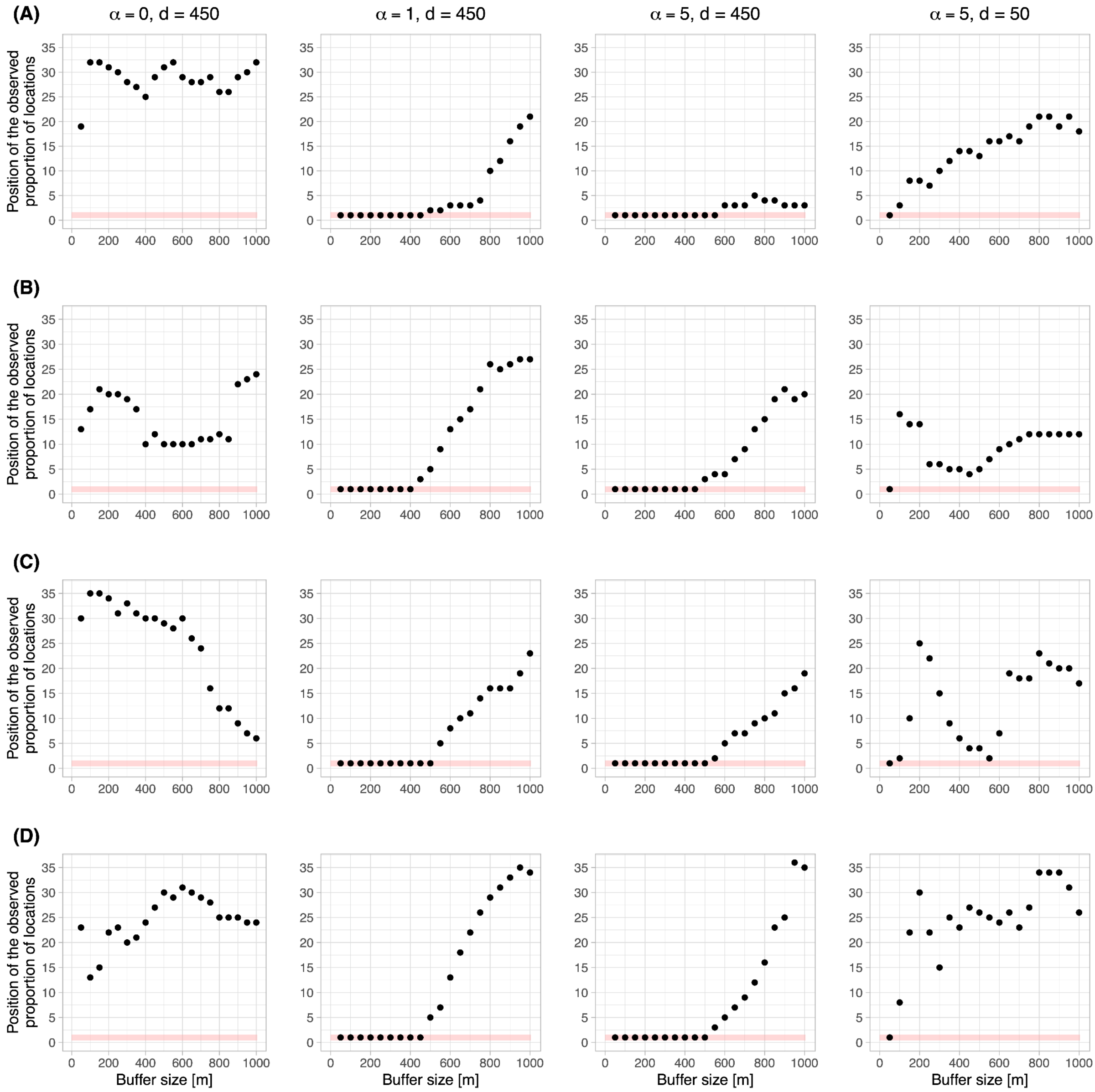


**Fig. S2. Rotation analysis results on simulated tracks under four avoidance behaviour scenarii**. Results are based on 1000 simulated tracks, each column representing a combination of turning angle ($\alpha$) and perceptual range ($d$). Rows refers to simulations done in different wind farms (hence with different wind turbine locations and configuration): La Baume (A), Montfrech (B), Mas de Naï (C) and Saint Affrique (D). Dots represent the position of the observed percentage of location, among the 36 estimated values ($y$-axis) during the rotational analysis, for a given circular buffer around wind turbines ($x$-axis). The red rectangle highlights buffers for which the amount of locations observed is significantly lower than randomly expected.

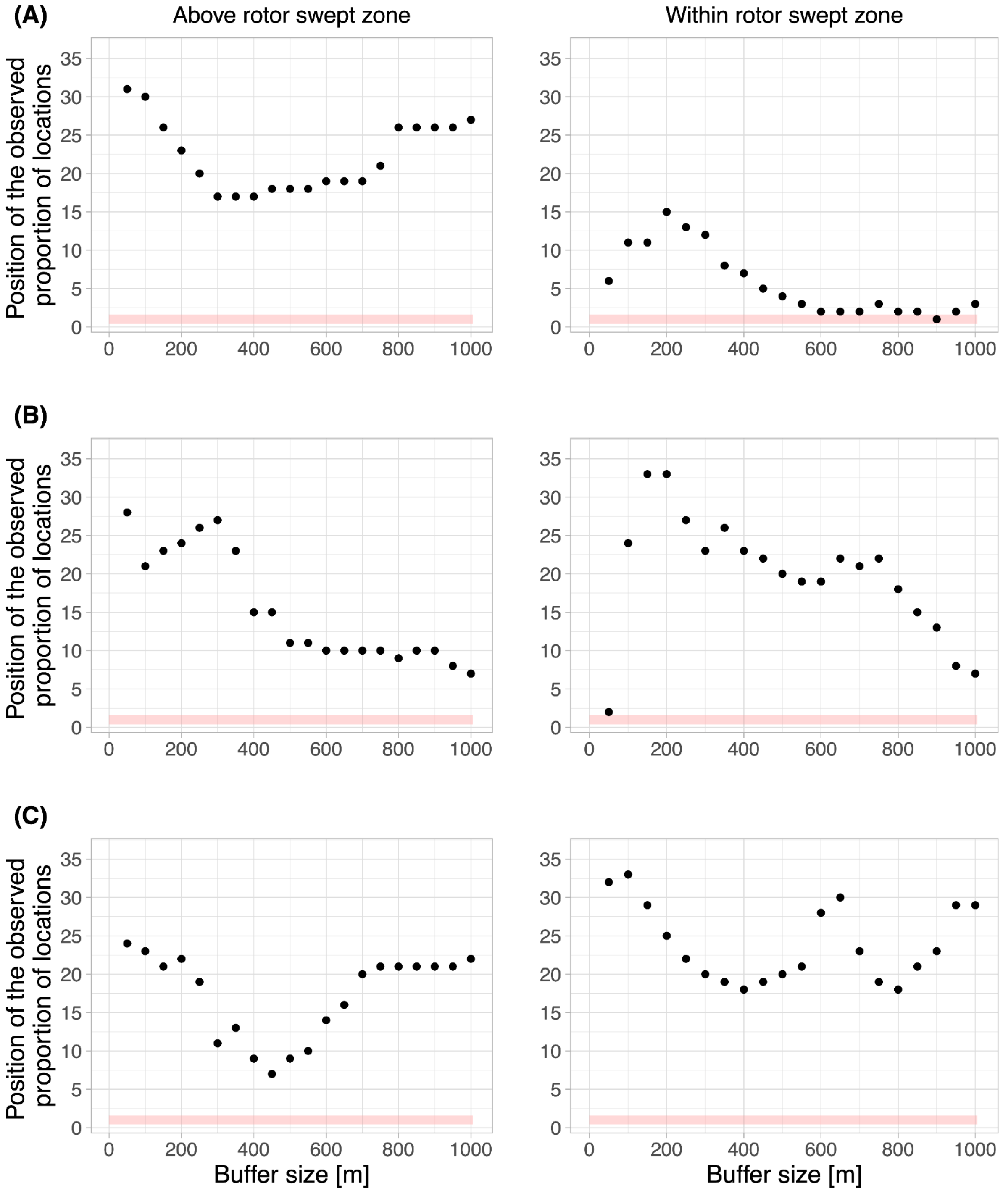


**Fig. S3. Rotation analysis results for vulture flights above (left) or within (right) the rotor swept zone of La Baume (A), Mas de Naï (B) and Saint Affrique (C) wind farms.** Dots represent the position of the observed percentage of location, among the 36 estimated values (*y*-axis) during the rotational analysis, for a given circular buffer around wind turbines (*x*-axis). The red rectangle highlights buffers for which the amount of locations observed is significantly lower than randomly expected.

**Table S1. Vulture-level data collection summary**. "Nb days of tracking" refers to the total number of days in which we had GPS data. The last four columns give the total number of flights that each individual did in each wind farm's geofence. These four wind farms were considered for meso-scale and micro-scale avoidance analysis (flights retained only if >10 consecutive GPS locations). Numbers in between brackets highlight the number of flights that entered the rotor swept zone. A "-" indicates that the individual did not cross the considered wind farm.

| Vulture names | Nb days of tracking | Nb flights in La Baume | Nb flights in Montfrech | Nb flights in Mas de Naï | Nb flights in Saint Affrique |
|---|---|---|---|---|---|
| Andouiller | 1095 | 15 (9) | - | 1 (0) | - |
| Bouffon | 1095 | 36 (19) | 1 (0) | 33 (6) | 1 (0) |
| Camembert | 681 | - | 16 (5) | - | - |
| Clarence | 1095 | 5 (2) | 1 (1) | 4 (3) | - |
| Claudine | 1095 | 46 (16) | 67 (21) | 3 (1) | 45 (15) |
| Cyrano | 1095 | 270 (127) | 30 (7) | 107 (46) | 91 (42) |
| EAC | 1095 | 311 (111) | 36 (14) | 80 (44) | 74 (33) |
| Goliath | 1095 | 9 (3) | 5 (2) | - | 1 (0) |
| Goulag | 1095 | 45 (24) | - | 88 (55) | - |
| Kilimandjaro | 341 | 1 (0) | 1 (0) | - | - |
| Millau | 1095 | 11 (3) | 4 (0) | 11 (5) | 1 (1) |
| Nemo | 1095 | 90 (53) | - | 17 (9) | 18 (8) |
| Obier | 870 | 3 (1) | - | - | - |
| Quaker | 1095 | 108 (60) | - | 4 (3) | - |
| Quintessence | 1095 | 527 (252) | 10 (5) | 5 (3) | 111 (55) |
| Quolibet | 1095 | 99 (61) | 1 (0) | 16 (7) | - |
| Rodin | 365 | 4 (1) | 7 (4) | 2 (1) | 7 (1) |
| Rostand | 1072 | 21 (6) | 4 (0) | 1 (0) | 6 (1) |
| Veto | 1095 | 2 (2) | - | - | - |
| Westminster | 1095 | - | 2 (1) | 1 (1) | - |
| Xixa | 1095 | 80 (31) | 21 (7) | 89 (50) | 98 (34) |
| Xorifiant | 1095 | 81 (37) | - | 17 (10) | 2 (0) |
| Yokohama | 1095 | 1 (0) | - | - | - |
| Yucca | 1095 | 9 (5) | - | 46 (20) | - |
| Kenjutsu | 1095 | 19 (13) | 1 (0) | 4 (1) | 1 (1) |

**Table S2. Wind farm-level data collection summary**. “Max” and “Min rotor height” columns refer to the distance (in metres) between the ground and the tip of wind turbine blades. “Nb tot flights” column gives the total number of flights (flights retained only if > to 10 consecutive GPS locations) detected within the wind farm geofence. “Percentage of flights crossing the rotor-swept zone” counts the number of flights that happened (at least partially) in the rotor swept zone (i.e. between Min rotor height and Max rotor height + 15 m). “Percentage of locations above the rotor swept zone” informs about the percentage of GPS locations that were above the rotor swept zone.

| Wind farm names | Nb turbines | Min rotor height | Max rotor height | Nb tot flights | Percentage of flights crossing the rotor swept zone | Percentage of locations above the rotor swept zone |
|---|---|---|---|---|---|---|
| La Baume - S1 | 6 | 25 | 125 | 1793 | 46.80 | 82.55 |
| Montfrech - S2 | 4 | 30 | 110 | 207 | 32.84 | 84.70 |
| Mas de Naï - S6 | 17 | 30 | 90 | 529 | 50.28 | 85.49 |
| Saint Affrique - S7 | 6 | 35 | 125 | 456 | 42.07 | 86.80 |
